# Social selection within aggregative multicellular development drives morphological evolution

**DOI:** 10.1101/2021.05.12.443771

**Authors:** Marco La Fortezza, Gregory J. Velicer

## Abstract

The evolution of developmental systems might be shaped by both historical differences in developmental features and social selection, among other factors. In aggregative multicellularity, development is itself a social process in which unicellular organisms cooperate in carrying out complex developmental programs. In some aggregative systems, development culminates in the construction of spore-packed fruiting bodies. Fruiting body development in myxobacteria often unfolds within genetically and behaviorally diverse conspecific cellular environments that can include social defection and warfare. Here we use the bacterium *Myxococcus xanthus* to test whether the character of the cellular environment during aggregative development shapes morphological evolution. We manipulated the cellular composition of *Myxococcus* development in an experiment in which evolving populations initiated from a single ancestor repeatedly co-developed with one of several non-evolving partners - a benign cooperator, one of three cheaters or one of three antagonists. Fruiting body morphology was found to diversify as a function of developmental partners, revealing adaptation specific to distinct cellular environments. Collectively, antagonistic partners selected for higher levels of robust fruiting body formation than did cheaters or the benign cooperator. Moreover, even small degrees of genetic divergence between the distinct cheater partners were sufficient to drive treatment-level morphological divergence. Co-developmental partners not only shaped mean trait evolution but also determined the magnitude and dynamics of stochastic morphological diversification and subsequent convergence. In sum, we find that even few genetic differences affecting developmental and social features can greatly impact the morphological evolution of multicellular bodies and experimentally demonstrate that microbial warfare can promote cooperation.

## Introduction

Multicellular developmental systems involving cell differentiation, complex cell-cell interactions and collective morphological output have originated independently many times in both zygotic (1) and aggregative modes (2, 3). Once originated, developmental systems have diversified remarkably at all organizational levels from interplay among selective, historical, and stochastic forces (4–6) to generate seemingly “endless forms” (7). Biotic selection shaping development occurs both across and within species, for example from predation (2, 8) and sexual selection (9, 10), respectively. Other social forces such as interference competition and cheating also have potential to shape the evolution of developmental systems but are less studied in this regard (9, 11–14). Developmental-system evolution might be shaped not only by external selective forces, but also by internal system features (4–7).

Diverse prokaryotic and eukaryotic microbes exhibit aggregative multicellular development (2, 15). In some microbial systems such as the predatory myxobacteria and dictyostelids, starvation induces conspecific unicellular organisms to cooperatively develop into multicellular fruiting bodies packed with stress-resistant spores that germinate upon encountering conducive conditions (16, 17). Like zygotic developmental systems, aggregative systems have complex genetic architectures and involve temporal cascades of cell-cell signaling resulting in cell-type differentiation. In the myxobacteria, fruiting body morphologies and their genetic foundations have diversified greatly (18, 19) (Figure 1A). Such morphological diversification may have been driven in part by variation in selective forces, whether abiotic, e.g. the physico-chemical character of soil surfaces (20), or biotic, e.g. the composition of prey communities (21) or social groups (22, 23). However, the roles of diverse potential selective forces relative to historical constraints and stochasticity in shaping extant morphological diversity are unknown.

**Figure 1.**
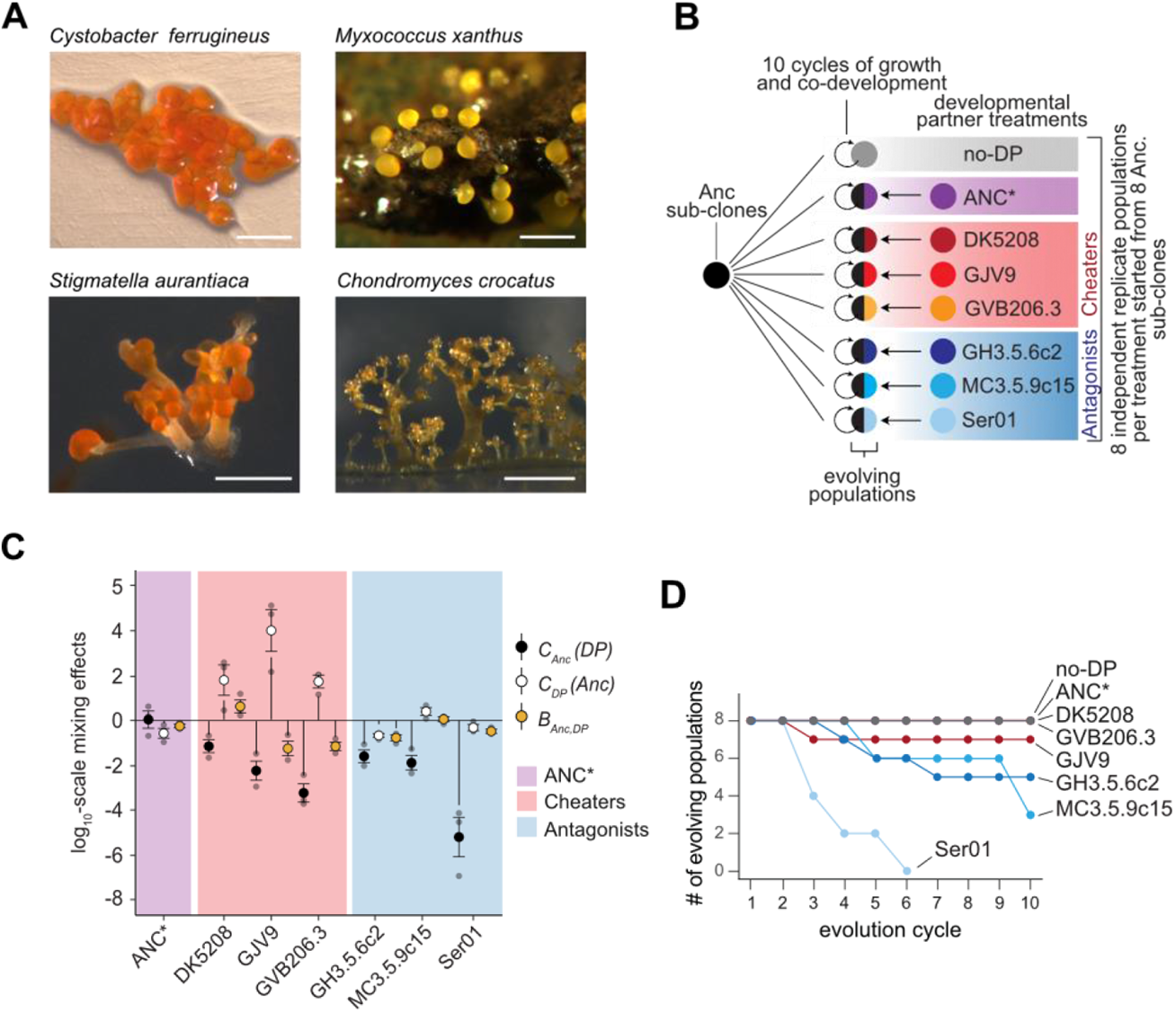
MyxoEE-7 tests whether distinct cellular environments during development shape morphological diversification in *M. xanthus*. **A**, Fruiting body morphologies across four Myxococcales species. Left to right: *C. ferrugineus* (Cb f17), *M. xanthus* fruiting bodies on soil, *S. aurantiaca* (Sg a32), and *C. crocatus* (DSM 14714T). (*M. xanthus* image from Michiel Vos, others from Garcia Ronald and Rolf Müller). Scale bars are ∼200 μm. **B**, MyxoEE-7 design summary. Black semi-circles represent Anc-derived evolving populations that repeatedly undergo co-development with a non-evolving partner (colored semicircles). The grey circle represents populations evolving with no developmental partner (no-DP). **C**, Effects of mixing the MyxoEE-7 *M. xanthus* ancestor Anc 1:1 with each non-evolving developmental partner (DP) on spore production by Anc [*C*_*Anc*_*(DP)*, black circles], DP [*C*_*DP*_*(Anc)*, withe circles] and the total group [*B*_*Anc,DP*_, tan circles] relative to monoculture spore production (see Material and Methods). Lines extending from zero indicate significant effects (one-sample t-tests, all p-values are listed in Table S1). A zero value indicates no effect of mixing (horizontal-solid line). See Figure S1A for pure-culture spore counts and absolute mix counts. Error bars, SEM (n = 3). **D**, Survival/extinction dynamics for all eight MyxoEE-7 treatments. The y-axis values indicate the number of independent replicate populations within each treatment defined by developmental partner (or none) at each evolution cycle.

The potential for genetic diversity among co-developing cells is an important feature of microbial aggregative development from onset to culmination. *Myxococcus* fruiting bodies emergent from natural soils are often composed of recently and locally diversified lineages and yet differentiated at the group level across small spatial scales (23, 24). Such fine-scale mosaic geographic structure of cell-group diversity appears to persist over evolutionarily significant periods (23). Thus, the evolution of aggregative development might be subject to group-level forms of historical contingency. The developmental evolution of a focal lineage may be contingent on not only its own genetic history (the standard conception of historical contingency (4, 5)), but also on the histories of other lineages with which it interacts, co-develops, and potentially co-evolves (22– 25). In the latter case, the relevant historical contingencies are also social selective forces. And even when fruiting bodies have little or no internal genetic diversity (23), the cellular environments of development generated by distinct founding genotypes might differ in their selective effects on new mutations and thus differentially shape future developmental evolution. Here we test whether the cellular environment of microbial aggregative development matters to the morphological evolution of fruiting bodies.

Interactions during development between distinct *M. xanthus* isolates sampled from the same fruiting body are often synergistic but sometimes antagonistic (22). Such groupmates often vary in monoculture spore production and include genotypes with severe developmental defects that can be socially complemented by developmentally-proficient genotypes from the same group (22-24, 26). Some defective genotypes “cheat” by exploiting proficient genotypes in mixed groups to gain a relative fitness advantage at spore production (27). Soil populations are structured in a fine-scale biogeographic mosaic of antagonistic conspecifics that combat one another upon encounter and that co-exist patchily due to positive frequency dependence of combat success (28). Such conspecific warfare is common in bacteria and is expected to promote various forms of microbial cooperation (28–32).

In light of potential natural interactions among myxobacterial conspecifics differing in social character, we designed an evolution experiment named MyxoEE-7 to address questions regarding effects of cellular diversity within developmental systems on their morphological evolution. Specifically, we ask whether morphological evolution of cooperative development can be differentially shaped by i) differences in the cellular composition of development *per se*, ii) major behavioral categories of developmental partners - benign cooperators, cheaters and antagonists (see Methods), and iii) small genetic differences (e.g. between distinct cheaters).

## Results

Starting with eight subclones of one ancestral genotype (Anc, Table S1), we established eight replicate populations in each of eight treatments to initiate MyxoEE-7 (Figure 1B, Figure S1A, Methods). Each evolutionary cycle consisted of growth in nutrient-rich liquid followed by development on starvation agar (Figure S1A). The cellular environment of development was manipulated in seven treatments by mixing Anc-derived populations 1:1 with a non-evolving developmental partner unique to each treatment at the onset of starvation (or no partner in one additional treatment) (Figure 1B). After development, spores were heat-selected and non-evolving partners were killed with antibiotic during growth (Figure S1A). We thus created eight distinct treatments of replicated developmental systems. Each system had one cellular component that would contribute descendants to the next developmental cycle and another cellular component that would not, with treatments initially differing only in the identity of their non-evolving developmental partner. We then tested whether the initially identical evolving components would morphologically diverge over time as a function of their non-evolving developmental partners. The three behavioral categories of non-evolving partners - benign cooperators, cheaters and antagonists - were hypothesized to impose different forms of social selection on Anc-derived populations during development and thus potentially differentially shape morphological evolution (Table S2). In the benign cooperator category, populations co-developed with the antibiotic-sensitive parent of the experimental ancestor Anc (ANC*, Figure 1B,C, Figure S1B and Table S1). In the cheater category, the three developmental partners were strains severely defective in monoculture sporulation that nonetheless outcompete Anc at sporulation in mixed groups (DK5208, GJV9, GVB206.3; Figure 1B,C, Figure S1B). In the antagonist category, evolving populations co-developed with one of three genomically distant natural isolates (GH3.5.6c2, MC3.5.9c15, and Ser01) that are highly proficient at fruiting body formation and sporulation in pure culture and greatly reduce Anc spore production (by ≥ 98%) in co-developmental groups (Figure 1B,C Figure S1B). Cheating and antagonistic partners are diverged from Anc by small (< 20 mutations) and large (> 100,000 mutations, core genome size ca. 9.1 Mb) degrees, respectively (Table S1). In an additional treatment with no developmental partner (no-DP, Figure 1B), the cellular composition of development was unmanipulated.

From an evo-devo perspective, the social character of aggregative development allowed us to manipulate the cellular environment of evolving developmental systems without starting evolution from multiple genetically distinct ancestors in a traditional historical-difference experiment (5). We thus test for an effect of the cellular context of developmental systems *per se* on morphological evolution. From a social evolution perspective, we examine how repeated interaction with conspecifics of radically different social character - friendly cooperators, closely related cheats, and distantly related antagonists - evolutionarily shapes the output of a cooperative process.

Seventeen (17) of the 64 initial MyxoEE-7 populations (27%) went extinct during the experiment, all but one of which co-developed with an antagonist (*p* < 0.0001, Fisher’s exact test). All eight populations partnered with the antagonist Ser01 went extinct by the sixth cycle (Figure 1D), an outcome explainable by the severe reductions of Anc spore production caused by co-development with Ser01 (Figure S1B). However, because Anc spore production was consistently ∼1,000-fold higher when mixed with GH3.5.6c2 or MC3.5.9c15 compared to mixes with Ser01 (Figure S1B), simple experimental variation in co-developmental outcomes is unlikely to explain the extinction events in GH3.5.9c2- and MC3.5.9c15-partnered populations. One possible explanation, among others, is the appearance of adaptive mutations that generate frequency-dependent sensitivity to antagonistic partners. In this scenario, low-frequency adaptive mutants receive social protection against antagonists from other high-frequency genotypes but increase in susceptibility to antagonists with increasing frequency. If such mutants reached sufficiently high frequency by the end of some growth phases, extinction events during subsequent development phases might result.

### The cellular character of aggregative development shapes morphological evolution

In comparison with the ancestor Anc, we quantified four morphological traits (see Methods) of all surviving populations in the absence of their developmental partner (if any) after four and ten cycles of experimental evolution (see Methods), thereby characterizing how intrinsic developmental phenotypes evolved in response to co-development with the various non-evolving partners. One trait - the number of fruiting bodies per plate - is a population-level phenotype whereas the three other traits are features of individual fruiting bodies - area, density, and density-heterogeneity (see Methods). We first integrate all traits in multivariate principal component analysis (PCA) to characterize overall morphological evolution and subsequently examine evolutionary patterns of individual traits.

The overall morphological analysis demonstrates that i) treatment-level sets of derived populations diverged from their ancestor and ii) the different non-evolving developmental partners caused their corresponding replicate-population sets to diversify at both the individual-treatment level and the treatment-category level (Figure 2B; Figure S2, Table S4). For example, already by cycle 4 most evolved treatments had diverged morphologically from Anc and the GJV9-partnered cheater treatment had collectively diverged from most other treatments (Figure 2B). By cycle 10, overall treatment-level divergence across the morphospace increased further still, with all three cheater-partner treatments showing morphological divergence from the two surviving antagonist-partner treatments (Figure 2B). Evidence of treatment divergence is additionally supported by hierarchical clustering analysis (see Methods) of overall morphological similarity, which indicates the evolution of three distinct morphological treatment clusters by cycle 10 (Figure S2C). Each cluster is composed of a pair of treatments that are morphologically more similar to one another than to any other treatment - a pair of cheater treatments (DK5208 and GJV9), the pair of surviving antagonist treatments and a pair formed by the benign cooperator ANC* treatment with the no-DP treatment (Figure S2C). Importantly, the morphological divergence observed at the treatment level demonstrates most fundamentally that populations underwent adaptation during the experiment and that some adaptation was specific to interaction with particular developmental partners. Since only the cellular environment of development differed systematically across treatments, only differential adaptation to distinct developmental partners can plausibly explain treatment-level divergence at heritable traits. The cellular character of aggregative developmental systems shaped the course of morphological evolution.

**Figure 2.**
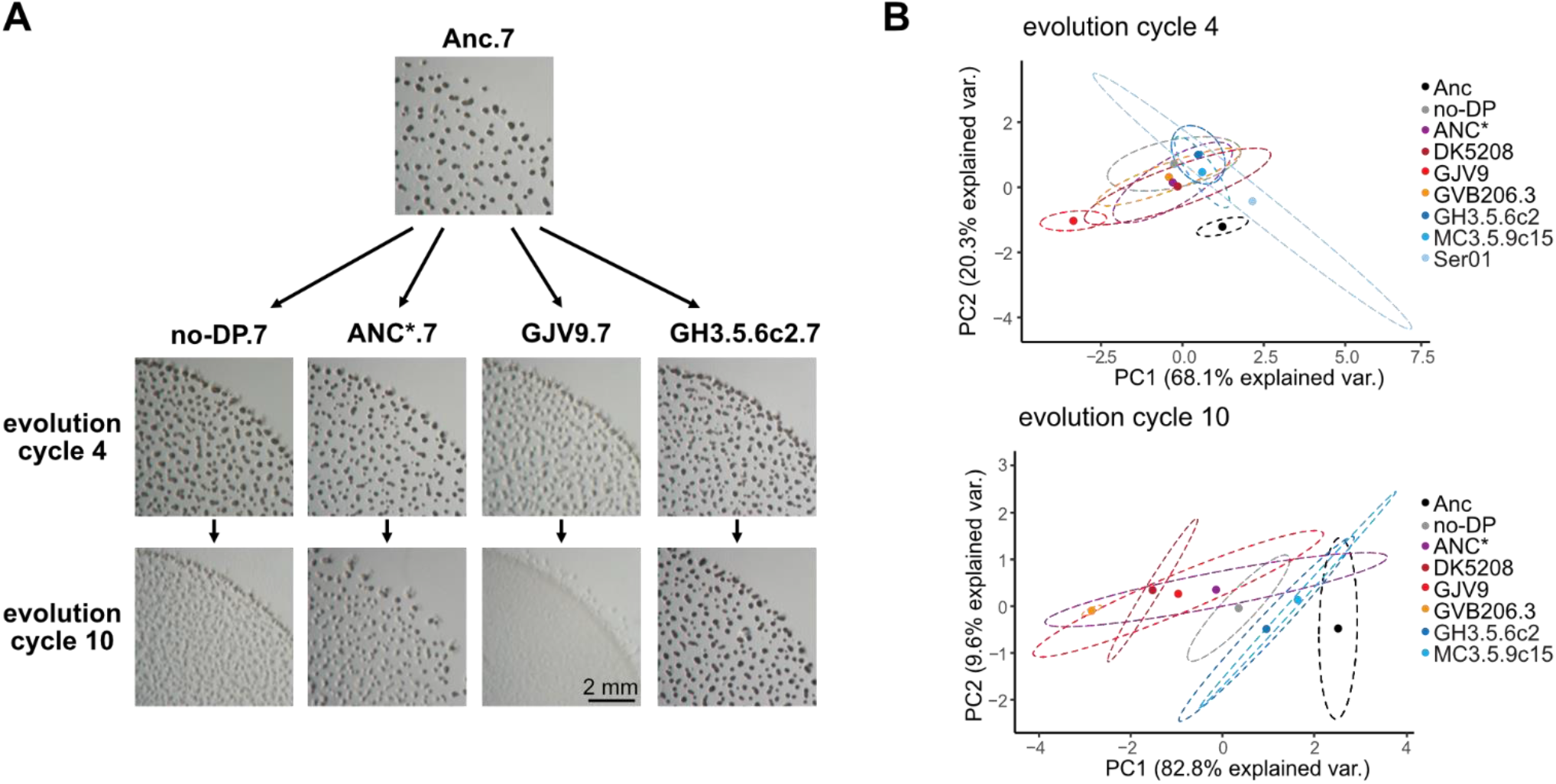
The cellular character of aggregative development shapes morphological diversification. **A**, Developmental phenotypes of the experimental ancestor and one evolved replicate population from four of the eight developmental-partner treatments after four and ten cycles of evolution. The evolved populations imaged here all descend from the same ancestral subclone (Anc.7). **B**, PCA-morphospace positions integrating four quantified developmental traits (Figure S2A) for the eight ancestral subclones (black circles and ellipses) used to initiate all evolution lines and for all treatment sets of evolved populations (colored circles and ellipses) after four and ten cycles of evolution. Circles represent PCA-morphospace centroids obtained from three biological-replicate assays surrounded by 95% confidence regions (dashed ellipses) (n = 3, Table S4). Percentage values on the x and y axes indicate the proportion of variation (‘var.’) explained by the first and second principal components, respectively (Figure S2B).

With the exception of fruiting body (FB) counts at cycle 4, average trait values across the entire experiment tended to decrease during evolution (Figure 3A), but this general trend masks rich variation in dynamics across, and in some cases within, developmental partner categories. Consistent with the divergent position of GJV9-partnered populations in PCA morphospace (Figure 2B), this cheater treatment showed unique evolutionary patterns at all four analyzed traits by cycle 4, decreasing rather than increasing in FB number and decreasing more than any other treatment at the other three traits (Figure S3A). Later, ANC* and the cheaters DK5208 and GVB206.3 drove large treatment-level decreases in FB number to levels near or below that of the GJV9-partnered treatment at cycle 10, which decreased only slightly, if at all, in the later cycles (Figure S3A). In one particularly striking comparison, GVB206.3-vs GJV9-partnered populations diverged greatly in FB number by cycle 4, with the GVB206.3 treatment making far more fruiting bodies, but then strongly reversed rank during later evolution (Figure S3A). The GVB206.3-partnered populations dropped to near zero FBs at cycle 10 while the GJV9-partnered populations continued to form hundreds of FBs on average (Figure S3A). Thus, one cheater (GJV9) drove a much more rapid decrease of FB formation while another (GVB206.3) ultimately drove a more complete loss. Given that ANC* and the three cheater partners are known or expected to differ from one another by only ∼7-30 mutations (Table S1), the unique trajectory of GJV9-partnered populations demonstrates that even small historical differences between developmental partners of focal evolving lineages can greatly impact morphological evolution of the latter. In social terminology, even small historical differences between group mates sharing a major behavioral characteristic - in this case cheating - can greatly impact how social phenotypes of focal lineages evolve.

**Figure 3.**
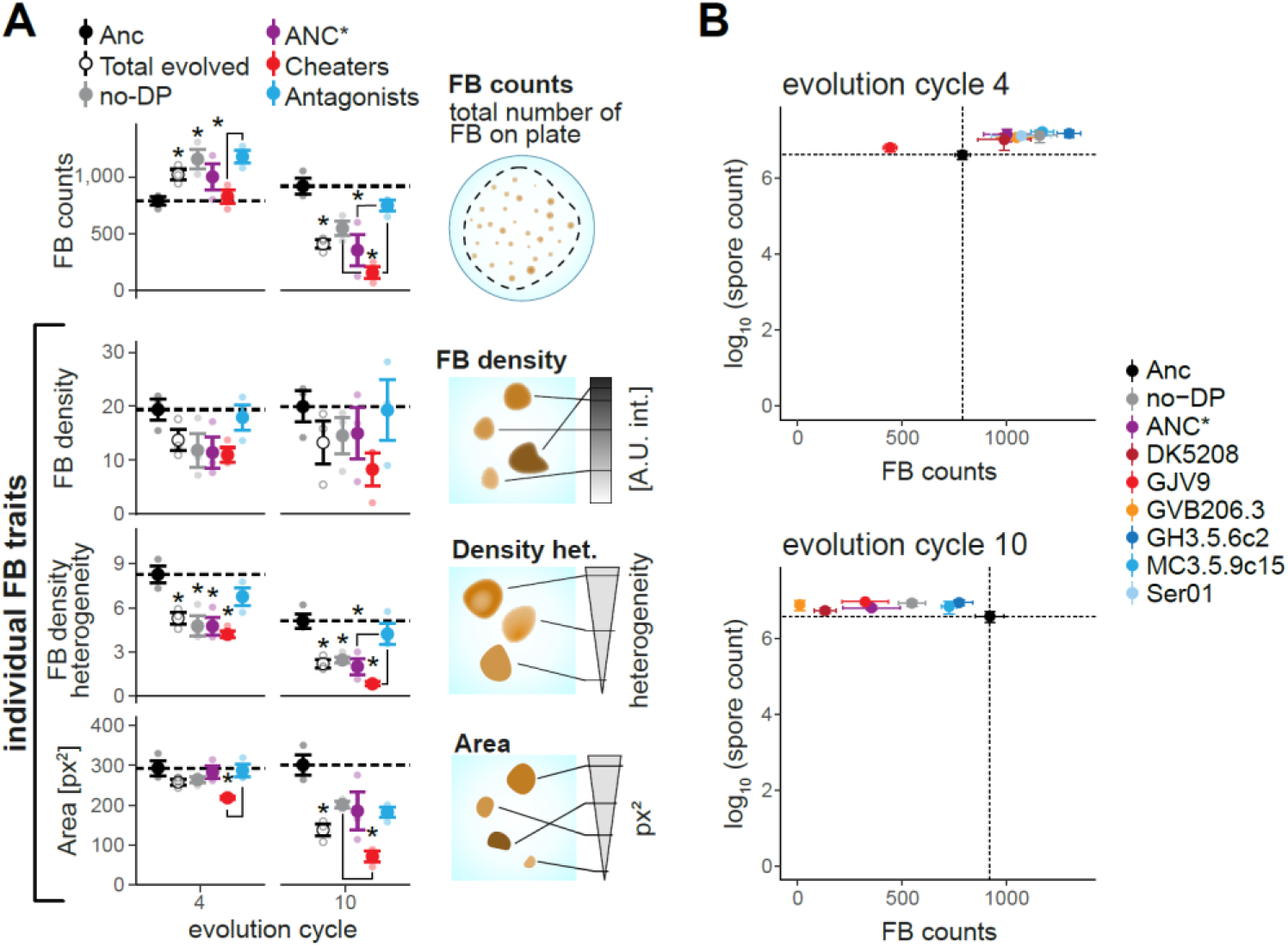
Social antagonists promote evolutionary maintenance of cooperative aggregative development. **A**, Phenotype means of the eight ancestral sub-clones (black circles), all evolved populations considered together (open circles), populations evolved with no partner (no-DP, grey circles), evolved populations partnered with ANC* (purple circles), all surviving populations partnered with a cheater (red circles) and all surviving populations partnered with an antagonist (blue circles) after four and ten evolutionary cycles. Circles represent cross-assay means (n = 3) of within-assay trait averages across all populations within each category. Stars represent significant differences relative to Ancestor and line connectors indicate significant differences between the respective categories of evolved populations (two-tailed t-tests were used to compare all evolved populations together (Total evolved) vs Anc, one-way ANOVA and subsequent two-tailed Tukey tests were used for all other contrasts, p values reported in Tables S5-7). **B**, Ancestral and treatment-level evolved spore production vs FB counts. In both a and b, ancestor means can differ between cycle 4 and cycle 10 plots because they were obtained independently for the two sets of separately-performed assays. In both a and b, error bars represent SEM.

### Distinct social interactant categories differentially affect morphological evolution

Morphological evolution varied not only between some individual treatments, but also among the three major partner categories defined by behavioral phenotypes - the benign parent (ANC*) of the experimental ancestor, cheaters, and antagonists (Figure 3A, Table S2). By cycle 4, surviving populations that co-developed with an antagonist collectively evolved higher FB counts than cheater-partnered populations (Figure 3A). By cycle 10, all categories decreased in FB counts from their respective cycle 4 values, but to different degrees. Specifically, ANC*- and cheater-partnered populations dropped to FB counts far below those of the Anc subclones, whereas antagonist-partnered populations continued to produce nearly as many fruiting bodies as Anc (Figure 3A). After ten cycles, ANC*- and cheater-partnered populations formed only 47% and 20% as many fruiting bodies as antagonist-partnered populations, respectively (Figure 3A). These large relative losses in FB numbers were not countered by relative gains at other traits indicative of developmental proficiency, particularly FB density or area. Indeed, FB-density and FB-area means for antagonist-partnered populations were either greater than or indistinguishable from means for the other categories at both evolved time points (Figure 3A, Figure S3A). These patterns collectively show that developmental interaction with antagonists selectively favored maintenance of robust fruiting body formation (e.g. Figure 2A) and thereby evolutionarily stabilized developmental cooperation.

The magnitude of morphological differentiation between treatments was not found to correlate with genomic divergence between respective developmental partners, as might be expected (5) (Table S1). Small genomic differences between GJV9 and the two other cheating partners (∼30 or fewer mutations, Table S1) caused greater divergence between their respective treatments at cycle 4 than did the much larger genomic difference (Table S1) between the antagonists GH3.5.6c2 and MC3.5.9c15 (Figure 2B; Figure S2C and 3A). Collectively, these outcomes indicate that small historical differences between developmental partners can strongly shape divergence of focal interactant lineages and suggest that some behavioral characteristics, e.g. antagonism, shared by even highly divergent partners can matter more for shaping developmental evolution than do large genomic differences.

### Sporulation is evolutionarily independent from fruiting body formation

Previous work suggested that evolutionary decreases in the ability to form many dense fruiting bodies in pure culture would generally be associated with large decreases in spore production (33), but this was not the case here. In all treatments except the GJV9-partnered treatment, both monoculture spore production and monoculture FB counts remained at or increased over the ancestral level by cycle 4 (Figure 3B, Figure S3B). However, among the GJV9-partnered populations at cycle 4, sporulation remained high while FB counts decreased greatly (∼40%) (Figure 3B, Figure S3), a pattern revealing evolutionary independence of prolific starvation-induced spore production from fruiting-body number. While *M. xanthus* can undergo a form of individualistic, chemical-induced sporulation by different pathways than starvation-induced sporulation (34, 35), pervasive evolutionary uncoupling of high levels of starvation-induced spore production from fruiting body formation is previously unknown. Further illustrating this independence, monoculture spore production remained at or above ancestral levels in all treatments at cycle 10 (Figure S3B) despite large decreases in FB counts, FB density and area in most treatments (Figure 3, Figure S3). Most strikingly, sporulation and FB formation became completely uncoupled among the GVB206.3-partnered populations, which made almost no FBs at cycle 10 while retaining ancestral levels of monoculture sporulation.

### Social selection limits the degree of stochastic morphological diversification

We additionally tested whether differences in adaptive-landscape topology imposed by distinct developmental partners differentially constrained morphological diversification among replicate populations within treatments (36, 37) (see Methods) - diversification which is caused by stochastic variation in mutational input. After four cycles, cheater-partnered population sets collectively evolved greater degrees of inter-population diversity within treatments than was present among the Anc sub-clones but antagonist-partnered populations did not (Figure 4). However, differential patterns of stochastic diversification were dynamic, shifting and even reversing over later cycles. Replicate populations with no partner and those partnered with ANC* began and continued increasingly diversifying by cycles 4 and 10, respectively. In contrast, antagonist-partner treatments underwent highly parallel evolution in the early cycles, only to later undergo a relative explosion of morphological diversification. In particular, the GH3.5.6c2 treatment ended MyxoEE-7 more diversified than all non-antagonist treatments. In yet further contrast, during early evolution the populations within cheater-partnered treatments underwent the greatest diversification but then later converged toward morphological similarity (Figure 4) while continuing to collectively diverge from their ancestor in trait means (Figure 3A). Strikingly, GVB206.3-partnered populations converged to near morphological identity (Figure 4). Late convergence within the cheater-partner treatments reflects trajectories of greatly decreased FB numbers, density, and area (Figure 3, Figure S3). In sum, cellular environments differentially determine the temporal dynamics and magnitude of stochastic developmental diversification and post-diversification convergence.

**Figure 4.**
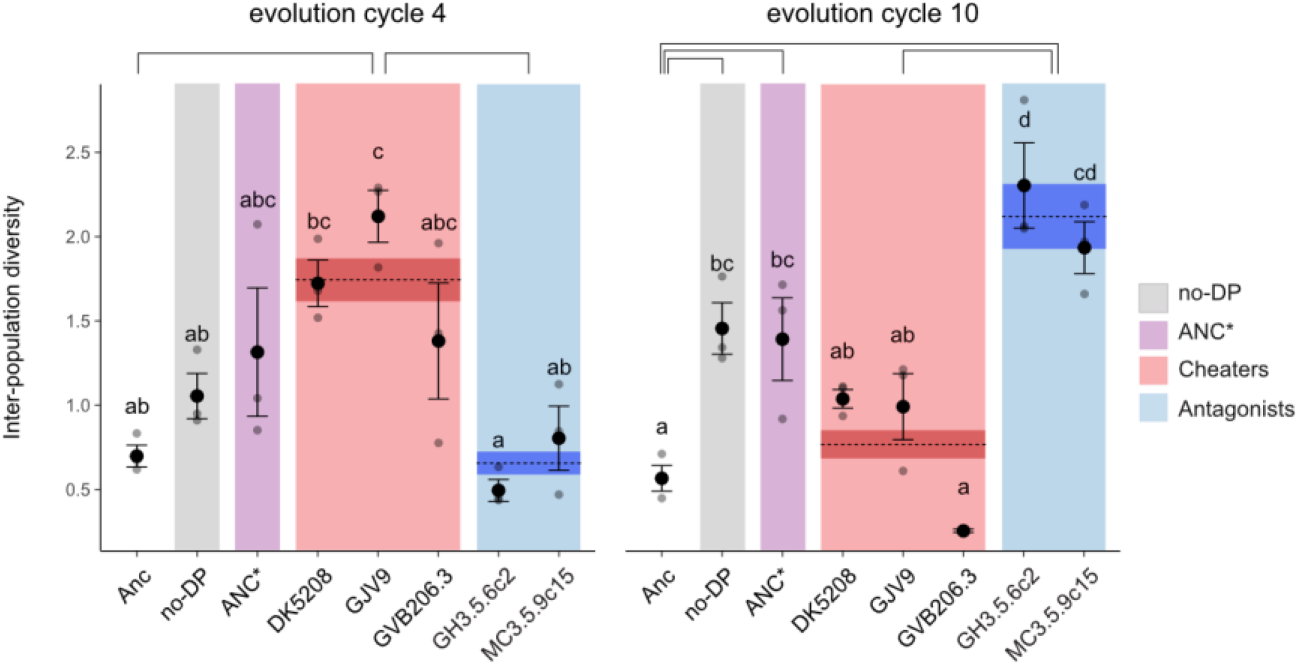
Developmental partners differentially determine patterns of stochastic morphological diversification and deterministic convergence. Quantification of morphological diversification (see Material and Methods) among the eight Anc subclones and among evolved populations within each cellular-environment treatment and treatment category after four and ten evolution cycles. Colored shading highlights treatment categories. Dashed lines indicate treatment-category means and surrounding darker shading indicates the corresponding SEM. Non-overlapping letters sets and connecting solid-black lines indicate significant differences between mean values of treatment and treatment categories, respectively (one-way ANOVA followed by two-tailed Tukey tests, Tables S12-14). The Ser01-antagonist treatment was not analyzed due to many early extinction events. Error bars, SEM (n = 3). Note that categories belonging to the same statistical class does not imply morphological similarity between the categories, but only indistinguishable degrees of within-category diversification. Compared categories might have identical diversification means while having very different trait means.

## Discussion

Social forces such as cooperation, cheating and interference competition have the potential to influence developmental processes and their evolutionary trajectories (9, 13, 14). In aggregative developmental systems, such social forces occur within the developmental process because conspecifics are the cells that carry out development. In this study, we manipulated the cellular composition of aggregative multicellular development in the bacterium *M. xanthus* during MyxoEE-7 (Figure 1B, Figure S1A, see Methods) and demonstrated that the character of cellular interactants within developmental systems profoundly shapes the future morphological evolution of those systems. Distinct developmental partners strongly influenced both the deterministic evolution (Figures 2, 3) and stochastic diversification (Figure 4) of developmental phenotypes. Moreover, such developmental-partner effects were observed at multiple levels of categorization, both between treatments in which distinct cheating developmental partners differed by only several mutations and between treatment sets that collectively differed in a major behavioral feature of the developmental partner, e.g. cheating vs. antagonism.

Biotic antagonisms strongly shape the evolution of developmental and social systems. Across species, predation drives much morphological evolution in prey (38, 39), selects for cooperative prey defense (40) and can promote the evolution of multicellularity (8, 41). Within species, interference competition shapes animal cooperation (42) and is proposed to evolutionarily promote some forms of microbial cooperation, whether directly or indirectly (28, 29, 31, 32, 43). The uniquely high retention of the intrinsic ability to form robust fruiting bodies among the antagonist-partnered populations of MyxoEE-7 experimentally demonstrates that interaction with hostile conspecifics can directly favor a microbial cooperative trait. Mechanistically, antagonists may have disfavored mutations decreasing costly production of extracellular adhesins necessary for robust fruiting body formation (44), mutations that may have been advantageous in non-antagonist treatments. Alternatively, antagonists may have selected for mutations altering motility behavior that generate preferential aggregation among Anc-derived cells into fruiting bodies with fewer antagonist cells.

Our results suggest that conspecific interference competition may, in concert with other forces (Table S2), promote maintenance of fruiting body formation in the wild and may have even contributed to its original emergence. In *Myxococcus*, success at contact-dependent warfare is positively frequency dependent (28). If a geographic mosaic of antagonist-type territories (22, 28, 31, 32, 45) existed prior to the origin of fruiting bodies, evolving the ability to aggregate and cohere prior to sporulation may have generated benefits of larger dispersal groups for success at conspecific combat upon dispersal.

More broadly, effects of conspecific interference competition on the evolution of microbial cooperation and multicellularity require further investigation. Such effects might be direct responses to antagonism-imposed selection - as occurred in this study - or indirect by-products resulting from toxin-based warfare contributing to high-relatedness population structures conducive to the evolution of cooperative traits (28, 31, 32). Evolution experiments can be conducted to test whether and how different mechanistic forms of warfare promote microbial behaviors involving potentially exploitable secretions, including biofilm formation and counter-weaponry. With both microbial and larger organisms, comparative effects on social evolution of interaction with antagonistic, cheating and benignly cooperative conspecifics, singly or in combination, can be examined in both monospecific and community contexts.

Our finding that distinct developmental partners can greatly impact morphological evolution in *M. xanthus* may help to understand the remarkable diversification of fruiting body morphologies observed across myxobacterial species (Figure 1A). Distinct cellular compositions of spatially clustered lineage set during natural cycles of development may have contributed significantly to this diversification by differentially impacting evolutionary trajectories (18, 19). Across a broader range of multicellular developmental systems, the cellular character of development may shape not only both deterministic and stochastic forms of morphological evolution, but also the evolution of phenotypic plasticity (46–48), developmental bias (49) and both the genetic basis (50) and evolvability of development (51, 52).

## Material and Methods

### Semantics

In the context of MyxoEE-7, **‘developmental system’** refers to the total population of cells on a given starvation plate, including, for the seven treatments with non-evolving developmental partners, both the evolving and non-evolving subpopulation components. For purposes of this study, **‘cheater’** refers to a strain participating in a social interaction between cells of two different genotypes during co-development in which the cheating interactant (e.g. GJV9) is defective at developmental spore production (partially or fully) in pure culture relative to the other interactant, which is highly proficient at spore production (e.g. Anc). In the cheating interaction, the developmentally-defective strain gains a relative fitness advantage over the proficient strain by social exploitation, which is defined as a gain in absolute fitness derived from social interaction compared to monoculture. In the context of this study, **‘antagonist’** refers to a strain proficient at a high level of spore production that strongly suppresses spore production by another strain of similarly high sporulation proficiency (e.g. Anc). Such antagonism between natural isolates during development is generally also manifested during vegetative growth, indicating that such negative interaction effects are generally due to interference-competition mechanisms rather than from behaviorally facultative defection from cooperation specific to mixed-group development. By **‘benign cooperator’**, we refer to a strain that is highly proficient at cooperative development but does not antagonize a focal genotype.

### Strains

All strains in this study except for three natural isolates derive from the lab reference strain DK1622 (53) (Table S1). DK5208 (aka LS523 (54)) is derived immediately from DK1622 through a transposon insertion in the developmental signaling gene *csgA* (55). The strain ANC*, aka GJV2 (54), is a spontaneous rifampicin-resistant mutant of GJV1, which in turn is a sub-cultured derivative of the original DK1622 strain that differs from the published sequence of DK1622 by five mutations (56). The strain Anc, aka GJV27, is an immediate derivative of ANC* (GJV2) with a plasmid encoding kanamycin resistance integrated at its Mx8 phage attachment site (54). GVB206.3 (27, 33) and GVB207.3 (33, 56, 57) (of which GJV9 is a spontaneous rifampicin-resistant mutant (54)) are clonal isolates from experimental populations that evolved for 1000 generations in liquid culture derived from GJV2 and GJV1, respectively (Table S1).

GVB207.3 differs from GJV1 by 14 mutations (54) and hence from ANC*/GJV2 by 15 known mutations. If a similar number of mutations separate GVB206.3 from ANC*/GJV2 (since GVB206.3 and GVB207.3 evolved for the same number of generations in the same evolution experiment), then GJV9 and GVB206.3 differ by ∼29 mutations (also accounting for the rifampicin-resistance mutation in GJV9). Given the five-mutation difference between the original DK1622 and GJV1 and assuming no additional mutations in DK5208 other than the transposon insertion that distinguishes the latter, then DK5208 differs from ANC* by seven known mutations and from GVB206.3 and GJV9 by 21 known mutations. GH3.5.6c2 and MC3.5.9c15 differ from the DK1622-derived strains by ∼200k mutations (23) while Ser01 differs from the DK1622-derived by >100k mutations (58) (Table S1).

### Experimental evolution - MyxoEE-7

Eight subclones of GJV27 (here referred to as ‘Anc’) were isolated (GJV27.1/Anc.1 - GJV27.8/Anc.8), stored frozen and used to separately establish eight independent replicate populations for each of eight evolutionary treatments (64 populations total, Figure 1b). Anc-subclones and the seven strains used as non-evolving partners were initially grown separately in CTT (59) liquid (32 °C, 300 rpm) (Figure S1B) from frozen stocks. Mid-log-phase cultures were centrifuged (5 min at 12000 rpm) and resuspended to ∼5 × 10^9^ cells/ml in TPM liquid buffer (59). Anc-subclone populations were mixed with the seven developmental partners in a 1:1 ratio and 50 μl of the mixed populations (∼2.5 × 10^8^ cells total, ∼1.25 × 10^8^ cells of each partner) were subsequently spotted on TPM starvation-agar plates (1.5% agar) (59). A 50-μl aliquot of each resuspended Anc-subclone culture was also spotted without mixing for the control treatment (no-DP) that lacked a non-evolving co-developmental partner.

Populations were incubated on TPM agar for 5 days at 32 °C and 90% rH and then heated at 50 °C for 3 h to kill non-spores. Cells were harvested by scalpel and transferred into CTT liquid with kanamycin, which killed the non-evolving partners (Figure S1C). Cultures of the Anc-derived populations were then grown (32 °C, 300 rpm) to mid-log phase. Cultures were deemed extinct if no growth was evident after 5 days. To eliminate residual kanamycin from the growth medium prior to mixing Anc-derived cultures with their respective non-evolving partner for the next developmental phase, the evolving cultures were centrifuged as above, washed by resuspension in 2 ml TPM liquid and centrifuged again prior to final resuspension at ∼5 × 10^9^ cells/ml in TPM liquid. For each subsequent cycle, cultures of the non-evolving partners were initiated from the same frozen stock, grown in CTT liquid, and processed as for the first cycle. Mixes of evolving populations with non-evolving partners and cultures of the no-DP populations were then processed as in the first cycle to initiate subsequent cycles of starvation on TPM agar 5 days in duration (occasionally 4). Samples of all evolving populations were stored frozen (−80 °C, 20% glycerol) at the end of each growth phase, including after a growth phase following the tenth cycle of development. After five cycles into the experiment, all populations were discarded due to universal media contamination and were subsequently re-initiated from stocks frozen after the prior cycle. To facilitate reference to the evolution experiment reported here as a whole, we name the experiment ‘MyxoEE-7’, with ‘MyxoEE’ meaning ‘Myxobacteria Evolution Experiment’ and ‘7’ indicating the temporal rank position of this first publication from MyxoEE-7 relative to the first publications from other MyxoEEs (60). Prior MyxoEEs from which studies have already been published are correspondingly named MyxoEE-1 (33), MyxoEE-2 (57, 61), MyxoEE-3 (25, 60, 62 - 65), MyxoEE-4 (66), MyxoEE-5 (54) and MyxoEE-6 (21).

### Population bottlenecks

All developmental cycles were initiated with evolving population sizes of ∼1.25 × 10^8^ cells for treatments with partners or ∼2.5 × 10^8^ cells for the no-partner treatment. These population sizes were then reduced during development to various degrees. Estimated average bottleneck population-size ranges for the first cycle of MyxoEE-7 were ∼5 × 10^6^ -1 × 10^7^ for Anc and ANC*-partnered populations, ∼9 × 10^3^ -6 × 10^6^ for the three treatments of cheater-partnered populations, ∼6 × 10^4^-1 × 10^5^ for the MC3.5.9c15- and GH3.5.6c2-partnered populations and, most severely, ∼100 individuals for the Ser01-partnered populations (Figure S1A). These bottlenecks are caused by the intended selective forces of heat selection and, in some cases, negative effects of the cheaters or antagonists. The bottleneck sizes are likely to have increased over evolutionary time due to adaptive evolution. Despite the severity of the Ser01-caused bottleneck, we proceeded with the Ser01 treatment under the expectation that at least some spores would be produced in each round of co-development and under the reasoning that adaptive mutants arising during the growth phase with increased resistance to Ser01 antagonism would be at a strong advantage and thereby potentially greatly increase the bottleneck population size.

### Development assays

Cells from frozen stocks of the eight ancestral subclones Anc.1-Anc.8 and all evolved populations from either cycle 4 and or cycle 10 were inoculated and grown in CTT liquid with kanamycin until they reached mid-exponential phase. We subsequently centrifuged (5 min at 12000 rpm) and resuspended cells in TPM liquid twice to a final density of ∼5 × 10^9^ cells/ml for all samples. We spotted 50 μl of the resuspension on plates (6 cm diameter) containing 5 ml of TPM hard (1.5%) agar and allowed the resulting spots to dry for 1 h in a laminar-flow hood. Once dried, all plates were incubated upside down at 32 °C, rH 90% for 5 days before microscopy images of each plate were taken. All developmental assays were replicated three times independently at separate times. Assays of cycle 4 and cycle 10 populations were performed separately, each including all Anc subclones in every replicate. Assays for morphological analysis were performed separately from spore-production assays.

### Imaging and trait quantification

Representative images of evolved populations (Figure 2A) were obtained with a Zeiss STEMI 2000 microscope and captured with a Nikon Coolpix S10 camera. Images for quantitative morphological analysis were taken using an Olympus SXZ16 microscope with an Olympus DP80 camera system. The image-acquisition protocol was identical across all samples (exposure time = 9.9 ms, lens = Olympus 0.5xPF, zoom = 1.25x, ISO = 200, illumination = BF built-in system). Images were processed with FIJI software (67) by duplication, conversion to 8-bit mode and segmented using the Triangle algorithm to identify individual fruiting bodies. Dust particles that appeared on plates while imaging were masked before the segmentation process. Segmentation analysis was performed over the image area covered by the spotted cell population. Segmented objects with an area value smaller than 20 px^2^ were automatically excluded while all larger objects were retained for image analysis. No maximum size limit or circularity restrictions were imposed. Regions obtained in this manner were then over-imposed on the original images and used to define the area in which phenotypes were subsequently quantified.

After fruiting body (FB) definition, four traits were used both individually and collectively to quantitatively characterize developmental morphology:

1. *FB number* (20, 68, 69) – total number of fruiting bodies generated by ∼2.5 × 10^8^ initial cells on a single developmental plate.
2. *FB density*– average grey-value intensity of pixels per FB. This measurement is expressed in arbitrary-unit grey intensity values. Grey-value intensity is expected to correlate with total cell and spore density within a fruiting body. This trait has also been previously used in referring to “mature” FBs (20).
3. *Density heterogeneity* – standard deviation of within-FB pixel-grey values (FB density). This parameter quantifies the heterogeneity of cell density within a developmental aggregate and it has been studied with different methods in earlier works (70, 71).
4. *FB area* (20, 68, 69, 72) – surface area occupied by each FB expressed in total pixel number within the defined FB border.

For further analysis we used the median values of FB density, density heterogeneity and area measurements per plate. Trait values for all eight Anc subclones were obtained in parallel with evolved populations during each biological assay replicate. For most analyses, trait values were averaged across replicate populations from the same evolutionary treatment (or across Anc subclones) for each replicate assay and the resulting treatment-level means were subsequently averaged across three independent biological assay replicates.

### Spore counts and mixing effect quantification

Spore counts were obtained as in ref. (73), except that 50 °C heat selection was imposed by heating TPM developmental plates for 3 h prior to harvest. Effects of mixing Anc and each non-evolving developmental partner reported in Figure 1B on each strain individually (*C*_*i*_(*j*)) and total mixed-group productivity (*B*_*ij*_) were also calculated as in ref. (73).

### Statistical analysis

All statistical analyses were conducted using the software R (v4.0.0) (74). Pairwise comparisons across treatments or treatment categories were done with one-way ANOVA followed by two-sided Tukey tests. Direct comparisons of single treatments to Anc were done with one-way ANOVA followed by two-tailed Dunnett tests. Individual pair differences between experimental groups were identified using two-tailed *t*-tests.

### Multivariate analysis

Morphological trait values were first averaged across all evolved population replicates from each evolutionary treatment and across the eight Anc-subclones Anc.1-Anc.8 in each biological assay replicate (three each for the cycle 4 and cycle 10 assays, which were performed separately). PCA was run on scaled values using the *stat::prcomp()* function (74) (Figure 2B, Figure S2A, 2B). Plots reporting the PCA results for the two main dimensions PC1 and PC2 were obtained using the *ggplot()* package (75) (Figure 2B). perMANOVA was used to test whether evolutionary identity (including both developmental-partner identity of evolved populations and ancestral identity of Anc) significantly structures the data dispersion. To do so, we first test the homoscedasticity of the data’s dispersion with the *vegan::betadisper()* (76) function followed by a post-hoc Tukey test run with *stats::TukeyHSD()*. After confirming homogeneity of variances, perMANOVA was performed with the *vegan::adonis()* (76) function (Table S4).

### Morphospace distance estimates

Pairwise Euclidean distances between all treatment centroids in the morphospace were calculated on all four multivariate dimensions for cycle 4 and cycle 10 datasets separately. Hierarchical clustering based on the average pairwise distances method were subsequently calculated using the *pvclust::pvclust()* (77) function for both the cycle 4 and cycle 10 distance matrices. Approximately unbiased (AU) *p*-values were obtained from the *pvclust()* (77) function and used to determine the formation of significant dendrogram nodes (α= 0.05, all values are reported directly on the Figure S2C).

### Inter-population morphological diversity

We used PCA output to test for differences in the degree of morphological diversification undergone by independent replicate populations within the same developmental-partner treatment. For the cycle 4 and cycle 10 data separately, PCA was performed on the scaled trait values of all evolved populations and Anc subclones for each biological-assay replicate, except within-treatment trait values were not averaged across population replicates as in the prior analysis. Within each treatment, we used data dispersion around the centroid to determine the level of inter-population morphological diversification. Dispersion of Euclidean distances was measured using the *vegan::betadisper()* (76) function and contrasted across all treatments and treatment clusters (Table S13 and S14).

## Author Contributions

M.L.F. and G.J.V. designed the research and wrote the manuscript; M.L.F. conducted the experiments, analyzed the data, and created the figures.

## Competing Interest Statement

Authors do not declare any conflict of interests.

## Data availability

All data, raw images and programming codes used for the analyses are available upon request to the corresponding author.

## Funding

This work was funded in part by an EMBO Long-Term Fellowship (ALTF 1208-2017) to M.L.F. and SNSF grant 31003B_6005 to G.J.V.

## Acknowledgments

The authors thank Garcia Ronald and Rolf Müller for providing images of myxobacterial fruiting bodies and Samay Pande, Marie Vasse, Mary Jane West-Eberhard and Sébastien Wielgoss for comments on the manuscript.

## Notes

### Competing Interest Statement

The authors have declared no competing interest.

