## Supplementary material for "Social selection within aggregative multicellular development drives morphological evolution": Figure S, Table S

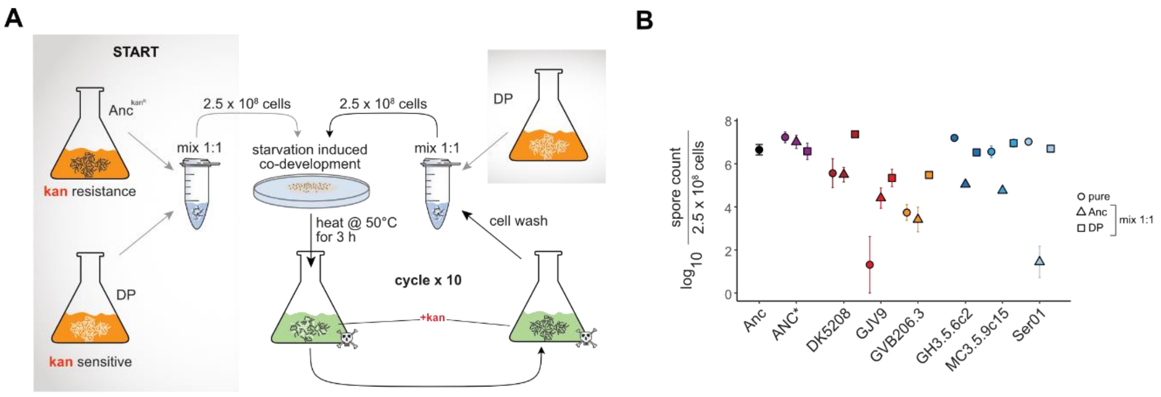


**Figure S1. Absolute spore production in co-development between Anc and non-evolving partners and MyxoEE-7 start and cycle summary.**

**A**, MyxoEE-7 start and cycle summary. Tan and green coloration indicates the absence or presence, respectively, of kanamycin in CTT liquid medium. **B**, Absolute spore production by Anc and all non-evolving developmental partners (DP) in pure culture (circles) and in 1:1 pairwise mixes of each partner with Anc (triangles - Anc with respective DP; squares - respective DP with Anc). Data points and error bars represent mean values and SEM, respectively (n = 3). Spore counts are standardized relative to ~2.5 x $10^8$ input cells, the total number of cells initiated in each developmental spot for both pure- and mixed-culture treatments. Pairwise mixes were initiated with ~1.25 x $10^8$ cells of each paired strain.

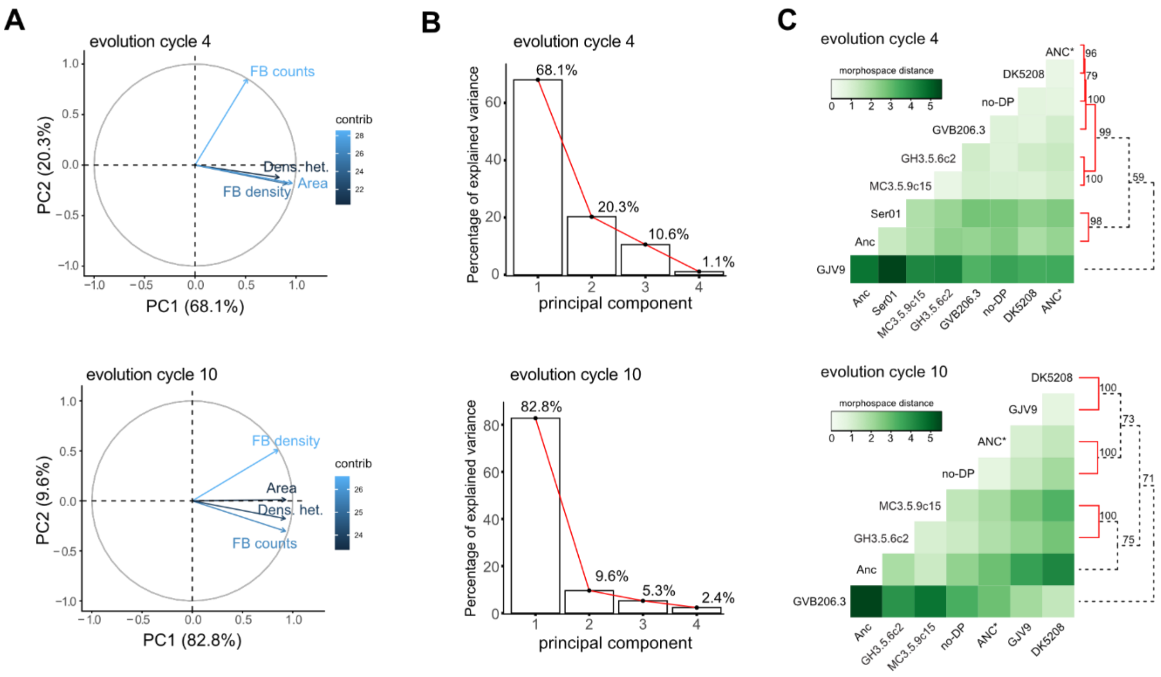


**Figure S2. PCA of phenotypic traits and analysis of treatment-level diversification.**

**A,** PCA eigenvectors of the four measured phenotypes colored from dark to light blue according to increasing contribution to shaping PCA morphospace at cycles 4 (top) and 10 (bottom). **B,** PCA Scree plots indicating variance percentages explained by the four principal components. **C,** Heatmaps of pairwise Euclidean distances between treatment-morphospace centroids at cycles 4 and 10. Cell darkness level reflects morphological distance between paired treatments. Dendrograms of hierarchical clustering across all pairwise inter-treatment distances in the morphospace are shown on the right side of each heatmap together with approximately unbiased (AU) *p*-values at each node. High AU *p* values (≥ 95) are considered to reflect significant morphological similarity between connected treatments or treatment sets (see Methods for details).

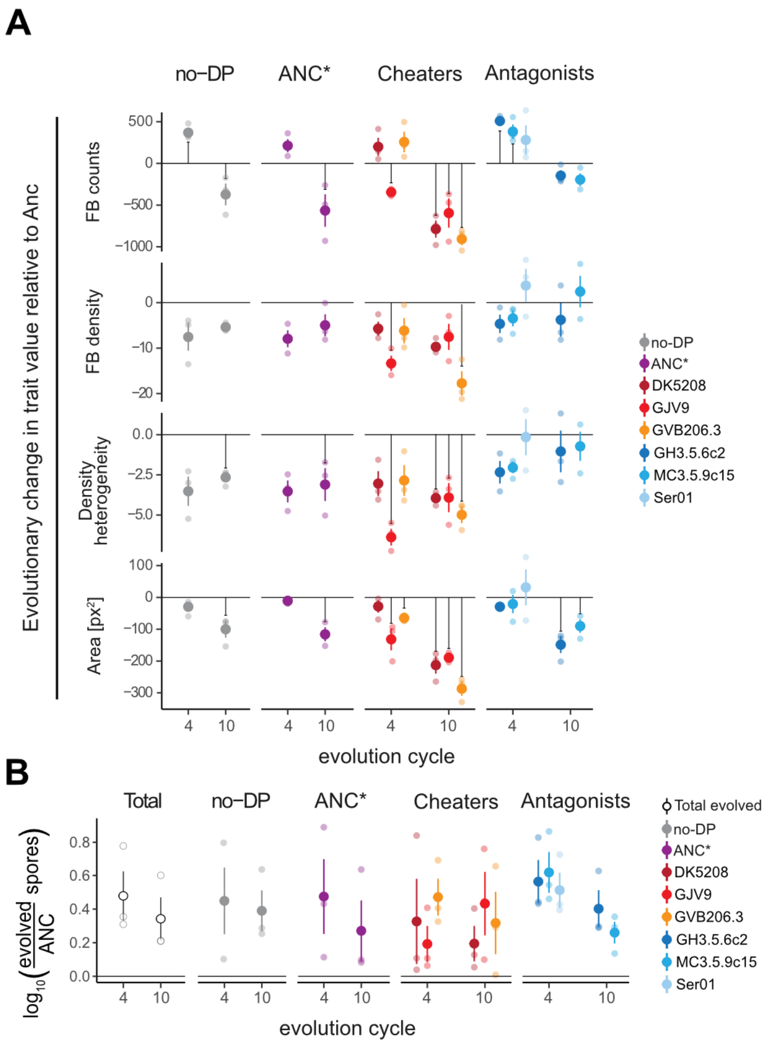


**Figure S3. Evolutionary changes in morphological trait values and spore production.**

**A,** Morphological trait means relative to Anc (horizontal-solid line set at zero) of evolved populations after repeated co-development with the benign ancestral cooperator (ANC*, purple circles), three cheaters (DK5208, dark-red circles; GJV9, bright-red circles; GVB206.3, orange circles) three antagonists (GH3.5.6c2, dark-blue circles; MC3.5.9c15, bright-blue circles; Ser01, light-blue circles) or no developmental partner (no-DP, grey circles) after four and ten evolutionary cycles. **B**, Mean values of spore counts relative to Anc (horizontal-black line set at zero) of all evolved populations combined (white circles) and by individual treatment (colored circles, symbols as in panel **A**) after four and ten evolutionary cycles. Circles and associated error bars represent means and SEM respectively ($n$ = 3) in both panels. Significant differences from Anc are indicated by vertical black lines (one-way ANOVAs followed by two-tailed Dunnett tests for evolved treatments or two-sided one-sample $t$-test for test of changed spore production across all evolved populations). $p$-values are reported in Table S6 and S8 (panel **A** tests) and Tables S9-S11 (panel **B** tests).

**Table S1**. Strains.

| Name | Description | Interaction with Anc | Known/estimated mutation differences from ANC* | Reference |
|---|---|---|---|---|
| DK1622 | ancestor of all lab strains used in this study, not used here | N/A | 6 | 53 |
| GJV1 | derived from DK1622, parent of GJV2 | N/A | 1 | 56 |
| ANC* (GJV2) | $rif^R$ mutant of GJV1 | neutral | - | 54 |
| DK5208 (LS523) | csgA mutant of DK1622, $oxytet^R$ | cheating | 7 | 54 |
| GVB206.3 | 1000-gen. lab descendant of GJV2, $rif^R$ | cheating | 14 | 27, 33 |
| GJV9 | 1000-gen. lab descendant of GJV1, $rif^R$ mutant of GVB207.3 | cheating | 16 | 33, 54, 57 |
| GH3.5.6c2 | natural isolate, Bloomington, Indiana | antagonism | ~ 200000 | 23 |
| MC3.5.9c15 | natural isolate, Bloomington, Indiana | antagonism | ~ 200000 | 23 |
| Ser01 | natural isolate, Serengeti National Park, Tanzania | antagonism | > 100000 | 58 |
| Anc (GJV27) | immediate ancestor all evolved populations in this study, mutant of GJV2 transformed with pDW79, $rif^R$, $kan^R$ | N/A | pDW79 plasmid | 54 |
| Anc.1-Anc.8 | subclones of Anc used to found the replicate populations of all eight MyxoEE-7 treatments | N/A | pDW79 plasmid | |

**Table S2**. Potential selective effects of MyxoEE-7 non-evolving co-developmental partners.

| Difference of non-evolving partner from Anc potentially imposing selection during co-development [a] | Difference from Anc known or expected | | |
|---|---|---|---|
| | ANC* | Cheater | Antagonist |
| Reduced intercellular signal production [b] | No | Yes | No |
| Production of molecules directly harmful to Anc cells [c] | No | No | Yes |
| Developmental signal-receptor incompatibilities [d] | No | No | no clear expectation |
| Motility behaviour [e] | No | Yes | Yes |
| Temporal dynamics of development [f] | No | Yes | Yes |
| Extracellular factors other than those noted above (e.g. adhesins, exopolysaccharides, siderophores, etc.) [g] | No | Yes | Yes |

**a.** Distinct non-evolving developmental partner categories were predicted to impose different forms of cellular selection on evolving lineages, potentially resulting in differential evolution of intrinsic morphological traits.

**b.** Developmentally defective cheaters produce less of one or more intercellular signals necessary for normal development than do proficient strains (27). The antagonist partners are not defective at developmental signaling because they are all proficient at both robust fruiting body formation and high levels of spore production (Figure S1B).

**c.** Severe contact-dependent interference competition is pervasive among natural isolates of *M. xanthus* (28), but such antagonism is unlikely to have evolved between the cheaters and Anc because only <20 mutations that evolved under laboratory conditions are known or expected to distinguish them (Table S1). Thus, we expect that direct cellular harm of Anc is a selective force unique to the antagonist category of treatments.

**d.** Evolution of signal-receptor incompatibilities for developmental signals between divergent M. xanthus lineages has not been documented but is a theoretical possibility for the antagonists, which differ from Anc by >100,000 polymorphisms (Table S1).

**e.** All developmental partners except ANC* differ from Anc in their population-level motility phenotypes under vegetative growth conditions, with the cheaters GJV9 and GVB206.3 exhibiting substantially slower swarming than Anc on hard agar and the cheater DK5208 and all three antagonists showing smaller differences in group swarming rate (unpublished data). Thus, differences in detailed cell-level motility behavior between Anc and the non-ANC* partners are likely.

**f.** Natural isolates often exhibit variation in the temporal dynamics of development (78).

**g.** The total sets and levels of extracellular secretions other than developmental signals, anti-competitor toxins and motility-related compounds are expected to differ between Anc and both cheaters and antagonists.

**Table S3.** Two-sided one-sample t-tests on mixing effects (μ = 0) (n = 3).

| DP mixed with Anc. [1:1] | Mixing effect parameter | t | df | p-val |
|---|---|---|---|---|
| ANC* | $C_{Anc}(DP)$ | 0.09 | 2 | 0.9366 |
| | $B_{ij}$ | -3.30 | 2 | 0.0810 ‡ |
| | $C_{(DP)}Anc$ | -2.60 | 2 | 0.1215 |
| DK5208 | $C_{Anc}(DP)$ | -4.06 | 2 | 0.0557 ‡ |
| | $B_{ij}$ | 2.12 | 2 | 0.1684 |
| | $C_{(DP)}Anc$ | 2.66 | 2 | 0.1169 |
| GJV9 | $C_{Anc}(DP)$ | -5.27 | 2 | 0.0341 * |
| | $B_{ij}$ | -3.87 | 2 | 0.0609 ‡ |
| | $C_{(DP)}Anc$ | 4.32 | 2 | 0.0496 * |
| GVB206.3 | $C_{Anc}(DP)$ | -8.00 | 2 | 0.0153 * |
| | $B_{ij}$ | -6.30 | 2 | 0.0243 * |
| | $C_{(DP)}Anc$ | 5.99 | 2 | 0.0268 * |
| GH3.5.6c2 | $C_{Anc}(DP)$ | -5.68 | 2 | 0.0296 * |
| | $B_{ij}$ | -5.95 | 2 | 0.0271 * |
| | $C_{(DP)}Anc$ | -7.58 | 2 | 0.0170 * |
| MC3.5.9c15 | $C_{Anc}(DP)$ | -5.85 | 2 | 0.0280 * |
| | $B_{ij}$ | 0.41 | 2 | 0.7239 |
| | $C_{(DP)}Anc$ | 2.28 | 2 | 0.1499 |
| Ser01 | $C_{Anc}(DP)$ | -5.94 | 2 | 0.0272 * |
| | $B_{ij}$ | -5.48 | 2 | 0.0317 * |
| | $C_{(DP)}Anc$ | -2.31 | 2 | 0.1468 |

Significant ($p < 0.05$) and nearly significant ($0.1 > p > 0.05$) values are highlighted with * and ‡ respectively.

**Table S4**. perMANOVA analysis for PCA by treatment and treatment categories.

| **contrast** | **cycle** | **df** | **F** | **R2** | **Pr(>F)** |
|---|---|---|---|---|---|
| treatment | 4 | 8 | 5.07 | 69.28% | <0.001 * |
| | 10 | 7 | 7.38 | 76.36% | <0.001 * |
| treatment cat. | 4 | 4 | 3.56 | 39.30% | 0.006 * |
| | 10 | 4 | 10.06 | 67.92% | <0.001 * |

Significant ($p < 0.05$) values are highlighted with ***.**

**Table S5**. Two-sided Welch two sample t-test on total morphological-trait values means relative to Ancestor (n = 3).

| cycle | trait | t | df | p-value |
|---|---|---|---|---|
| 4 | FB counts | 3.98 | 3.7 | 0.0187 * |
| | FB density | -2.00 | 4.0 | 0.1156 |
| | Dens. het. | -4.28 | 3.5 | 0.0169 * |
| | Area | -1.75 | 2.6 | 0.1944 |
| 10 | FB counts | -6.50 | 3.1 | 0.0067 * |
| | FB density | -1.35 | 3.6 | 0.2543 |
| | Dens. het. | -5.08 | 3.1 | 0.0133 * |
| | Area | -5.41 | 3.3 | 0.0099 * |

Significant ($p < 0.05$) values are highlighted with *.

**Table S6.** One-way ANOVAs on single morphological traits across treatment categories and single treatments (n = 3).

| contrast | cycle | trait | df | F | Pr(>F) |
|---|---|---|---|---|---|
| treatment. categ. | 4 | FB counts | 4 | 5.94 | 0.0103 * |
| | | FB density | 4 | 2.70 | 0.0925 ‡ |
| | | Dens. het. | 4 | 9.10 | 0.0023 * |
| | | Area | 4 | 4.77 | 0.0206 * |
| | 10 | FB counts | 4 | 13.91 | 0.0004 * |
| | | FB density | 4 | 1.32 | 0.3320 |
| | | Dens. het. | 4 | 13.54 | 0.0005 * |
| | | Area | 4 | 9.88 | 0.0017 * |
| treatment | 4 | FB counts | 8 | 7.70 | 0.0002 * |
| | | FB density | 8 | 3.07 | 0.023 * |
| | | Dens. het. | 8 | 6.33 | 0.0006 * |
| | | Area | 8 | 4.84 | 0.0026 * |
| | 10 | FB counts | 7 | 17.04 | <0.0001 * |
| | | FB density | 7 | 1.99 | 0.12 |
| | | Dens. het. | 7 | 12.65 | <0.0001 * |
| | | Area | 7 | 14.02 | <0.0001 * |

Significant ($p < 0.05$) and nearly significant ($0.1 > p > 0.05$) values are highlighted with * and ‡ respectively.

**Table S7.** Two-sided Tukey test on morphological trait-mean differences between treatment categories (n = 3).

| | | cycle 4 | | cycle 10 | |
|---|---|---|---|---|---|
| **trait** | **contrast** | **t** | **adj. p-val.** | **t** | **adj. p-val.** |
| FB counts | Anc. ~ ANC* | -2.03 | 0.3206 | 4.90 | 0.0044 * |
| | Antag. ~ ANC* | 1.69 | 0.4826 | 3.42 | 0.0407 * |
| | Cheat. ~ ANC* | -1.67 | 0.4910 | -1.72 | 0.4641 |
| | no-DP ~ ANC* | 1.49 | 0.5929 | 1.67 | 0.4894 |
| | Antag. ~ Anc. | 3.71 | 0.0260 * | -1.58 | 0.5958 |
| | Cheat. ~ Anc. | 0.36 | 0.9960 | -6.62 | < 0.0001 * |
| | no-DP ~ Anc. | 3.51 | 0.0355 * | -3.23 | 0.0550 ‡ |
| | Cheat. ~ Antag. | -3.36 | 0.0453 * | -5.14 | 0.0032 * |
| | no-DP ~ Antag. | -0.20 | 0.9996 | -1.75 | 0.4488 |
| | no-DP ~ Cheat. | 3.15 | 0.0618 ‡ | 3.39 | 0.0426 * |
| FB density | Anc. ~ ANC* | 2.30 | 0.2220 | 0.86 | 0.9050 |
| | Antag. ~ ANC* | 1.88 | 0.3850 | 0.74 | 0.9410 |
| | Cheat. ~ ANC* | -0.14 | 1.0000 | -1.15 | 0.7730 |
| | no-DP ~ ANC* | 0.12 | 1.0000 | -0.07 | 1.0000 |
| | Antag. ~ Anc. | -0.42 | 0.9920 | -0.12 | 1.0000 |
| | Cheat. ~ Anc. | -2.43 | 0.1830 | -2.02 | 0.3240 |
| | no-DP ~ Anc. | -2.18 | 0.2600 | -0.93 | 0.8780 |
| | Cheat. ~ Antag. | -2.01 | 0.3250 | -1.90 | 0.3750 |
| | no-DP ~ Antag. | -1.76 | 0.4410 | -0.82 | 0.9200 |
| | no-DP ~ Cheat. | 0.25 | 0.9990 | 1.09 | 0.8100 |
| Dens. het. | Anc. ~ ANC* | 4.37 | 0.0094 * | 4.67 | 0.0060 * |
| | Antag. ~ ANC* | 2.50 | 0.1669 | 3.35 | 0.0460 * |
| | Cheat. ~ ANC* | -0.70 | 0.9525 | -1.78 | 0.4350 |
| | no-DP ~ ANC* | 0.01 | 1.0000 | 0.68 | 0.9572 |
| | Antag. ~ Anc. | -1.86 | 0.3878 | -1.33 | 0.6820 |
| | Cheat. ~ Anc. | -5.07 | 0.0034 * | -6.45 | <0.0001 * |
| | no-DP ~ Anc. | -4.36 | 0.0096 * | -4.00 | 0.0168 * |
| | Cheat. ~ Antag. | -3.20 | 0.057 ‡ | -5.12 | 0.0032 * |
| | no-DP ~ Antag. | -2.49 | 0.1694 | -2.67 | 0.1297 |
| | no-DP ~ Cheat. | 0.71 | 0.9500 | 2.45 | 0.1781 |
| Area | Anc. ~ ANC* | 0.55 | 0.9801 | 3.16 | 0.0613 ‡ |
| | Antag. ~ ANC* | 0.24 | 0.9992 | -0.09 | 1.0000 |
| | Cheat. ~ ANC* | -3.26 | 0.0521 ‡ | -3.10 | 0.0671 ‡ |
| | no-DP ~ ANC* | -0.95 | 0.8717 | 0.43 | 0.9914 |
| | Antag. ~ Anc. | -0.31 | 0.9976 | -3.25 | 0.0530 ‡ |
| | Cheat. ~ Anc. | -3.81 | 0.0255 * | -6.26 | <0.0001 * |
| | no-DP ~ Anc. | -1.49 | 0.5882 | -2.73 | 0.1191 |
| | Cheat. ~ Antag. | -3.50 | 0.0362 * | -3.01 | 0.0776 ‡ |
| | no-DP ~ Antag. | -1.18 | 0.7608 | 0.53 | 0.9822 |
| | no-DP ~ Cheat. | 2.31 | 0.2171 | 3.54 | 0.0342 * |

Significant ($p < 0.05$) and nearly significant ($0.1 > p > 0.05$) values are highlighted with * and ‡ respectively.

**Table S8.** Two-sided Dunnett test on single morphological trait-mean differences across treatments relative to Ancestor (n = 3).

| | | cycle 4 | | cycle 10 | |
|---|---|---|---|---|---|
| **trait** | **treatment** | **t** | **adj. p-val.** | **t** | **adj. p-val.** |
| FB counts | no-DP | 2.89 | 0.0551 ‡ | -3.38 | 0.0204 * |
| | ANC* | 1.67 | 0.4525 | -5.13 | <0.0001 * |
| | DK5208 | 1.56 | 0.5196 | -7.15 | <0.0001 * |
| | GJV9 | -2.70 | 0.0800 ‡ | -5.41 | <0.0001 * |
| | GVB206.3 | 2.02 | 0.2691 | -8.24 | <0.0001 * |
| | GH3.5.6c2 | 3.98 | 0.0055 * | -1.33 | 0.6417 |
| | MC3.5.9c15 | 2.98 | 0.0457 * | -1.77 | 0.3718 |
| | Sero1 | 2.20 | 0.1996 | | |
| FB density | no-DP | -1.91 | 0.3177 | -0.87 | 0.9130 |
| | ANC* | -2.01 | 0.2706 | -0.80 | 0.9386 |
| | DK5208 | -1.45 | 0.5949 | -1.57 | 0.4856 |
| | GJV9 | -3.38 | 0.0199 * | -1.22 | 0.7191 |
| | GVB206.3 | -1.56 | 0.5195 | -2.87 | 0.0556 ‡ |
| | GH3.5.6c2 | -1.18 | 0.7745 | -0.61 | 0.9838 |
| | MC3.5.9c15 | -0.88 | 0.9320 | 0.39 | 0.9988 |
| | Sero1 | 0.95 | 0.8996 | | |
| Dens. het. | no-DP | -3.27 | 0.0254 * | -3.75 | 0.0096 * |
| | ANC* | -3.28 | 0.0249 * | -4.38 | 0.0026 * |
| | DK5208 | -2.82 | 0.0623 ‡ | -5.58 | <0.0001 * |
| | GJV9 | -5.92 | <0.0001 * | -5.53 | <0.0001 * |
| | GVB206.3 | -2.65 | 0.0883 ‡ | -7.04 | <0.0001 * |
| | GH3.5.6c2 | -2.16 | 0.2089 | -1.47 | 0.5503 |
| | MC3.5.9c15 | -1.90 | 0.3225 | -1.02 | 0.8398 |
| | Sero1 | -0.14 | 1.0000 | | |
| Area | no-DP | -1.00 | 0.8775 | -3.03 | 0.0413 * |
| | ANC* | -0.37 | 0.9997 | -3.51 | 0.0158 * |
| | DK5208 | -0.96 | 0.8956 | -6.44 | <0.0001 * |
| | GJV9 | -4.49 | 0.0018 * | -5.73 | <0.0001 * |
| | GVB206.3 | -2.21 | 0.1963 | -8.68 | <0.0001 * |
| | GH3.5.6c2 | -1.00 | 0.8796 | -4.50 | 0.0021 * |
| | MC3.5.9c15 | -0.71 | 0.9771 | -2.73 | 0.0728 ‡ |
| | Sero1 | 1.08 | 0.8363 | | |

Significant ($p < 0.05$) and nearly significant ($0.1 > p > 0.05$) values are highlighted with * and ‡ respectively.

**Table S9.** Two-sided one-sample t-test on spore counts (μ = 0) (n = 3).

| cycle | t | df | p-val |
|---|---|---|---|
| 4 | 2.73 | 2 | 0.1119 |
| 10 | 3.12 | 2 | 0.0894 ‡ |

Nearly significant (0.1 > $p$ > 0.05) values are highlighted with ‡.

**Table S10.** One-way ANOVA on spore counts by treatment (n = 3).

| cycle | df | F | Pr(>F) |
|---|---|---|---|
| 4 | 8 | 1.39 | 0.268 |
| 10 | 7 | 1.40 | 0.27 |

**Table S11.** Two-sided Dunnett test on spore counts mean differences relative to Ancestor (n = 3).

| | cycle 4 | | cycle 10 | |
|---|---|---|---|---|
| treatment | t | adj. p-val. | t | adj. p-val. |
| no-DP | 1.86 | 0.3420 | 2.13 | 0.2100 |
| ANC* | 2.09 | 0.2400 | 1.17 | 0.7490 |
| DK5208 | 1.25 | 0.7290 | 0.70 | 0.9670 |
| GJV9 | 0.46 | 0.9980 | 2.45 | 0.1220 |
| GVB206.3 | 1.55 | 0.5240 | 1.76 | 0.3750 |
| GH3.5.6c2 | 2.21 | 0.1950 | 2.24 | 0.1760 |
| MC3.5.9c15 | 2.54 | 0.1080 | 1.40 | 0.5990 |
| Ser01 | 1.81 | 0.3690 | | |

**Table S12**. One-way ANOVA on inter-population diversity across treatment categories and single treatments (n = 3).

| contrast | cycle | df | F | Pr(>F) |
|---|---|---|---|---|
| treatment cat. | 4 | 4 | 5.47 | 0.0135 * |
| | 10 | 4 | 14.27 | 0.0004 * |
| treatment | 4 | 7 | 6.49 | 0.0010 * |
| | 10 | 7 | 16.84 | <0.0001 * |

Significant ($p$ < 0.05) values are highlighted with *.

**Table S13**. Two-sided Tukey test on inter-population diversity mean differences across treatments (n = 3).

| | cycle 4 | | cycle 10 | |
|---|---|---|---|---|
| **contrast** | **t** | **adj. p-val.** | **t** | **adj. p-val.** |
| no-DP~Anc. | 1.17 | 0.9288 | 3.81 | 0.0257 * |
| ANC*~Anc. | 2.03 | 0.4939 | 3.54 | 0.0436 * |
| GH3.5.6c2 ~Anc. | -0.67 | 0.9967 | 7.44 | <0.0001 * |
| MC3.5.9c15~Anc. | 0.35 | 1.0000 | 5.86 | <0.0001 * |
| DK5208~Anc. | 3.37 | 0.0592 ‡ | 2.02 | 0.4997 |
| GJV9~Anc. | 4.68 | 0.0047 * | 1.82 | 0.6158 |
| GVB206.3~Anc. | 2.24 | 0.3759 | -1.34 | 0.8723 |
| ANC*~no-DP | 0.86 | 0.9862 | -0.27 | 0.99999 |
| GH3.5.6c2~no-DP | -1.84 | 0.6030 | 3.63 | 0.0362 * |
| MC3.5.9c15~no-DP | -0.83 | 0.9889 | 2.05 | 0.4801 |
| DK5208~no-DP | 2.20 | 0.4004 | -1.79 | 0.6783 |
| GJV9-no~DP | 3.51 | 0.0455 * | -1.99 | 0.5176 |
| GVB206.3~no-DP | -1.08 | 0.9533 | -5.14 | 0.0019 * |
| GH3.5.6c2~ANC* | -2.70 | 0.1917 | 3.91 | 0.0211 |
| MC3.5.9c15~ANC* | -1.68 | 0.6982 | 2.32 | 0.3382 |
| DK5208~ANC* | 1.34 | 0.8692 | -1.52 | 0.7876 |
| GJV9~ANC* | 2.65 | 0.2066 | -1.72 | 0.6783 |
| GVB206.3~ANC* | 0.21 | 1.0000 | -4.87 | 0.00330 * |
| MC3.5.9c15~GH3.5.6c2 | 1.01 | 0.9644 | -1.58 | 0.7543 |
| DK5208~GH3.5.6c2 | 4.04 | 0.0163 * | -5.42 | 0.0012 * |
| GJV9~GH3.5.6c2 | 5.35 | 0.0014 * | -5.62 | <0.0001 * |
| GVB206.3~GH3.5.6c2 | 2.92 | 0.1330 | -8.78 | <0.0001 * |
| DK5208~MC3.5.9c15 | 3.02 | 0.1109 | -3.84 | 0.0240 * |
| GJV9~MC3.5.9c15 | 4.33 | 0.0095 * | -4.04 | 0.0164 * |
| GVB206.3~MC3.5.9c15 | 1.90 | 0.5696 | -7.19 | <0.0001 * |
| GJV9~DK5208 | 1.31 | 0.8822 | -0.20 | 1.0000 |
| GVB206.3~DK5208 | -1.13 | 0.9416 | -3.35 | 0.0615 ‡ |
| GVB206.3~GJV9 | -2.44 | 0.2888 | -3.15 | 0.0879 ‡ |

Significant ($p < 0.05$) and nearly significant ($0.1 > p > 0.05$) values are highlighted with * and ‡ respectively.

**Table S14**. Two-sided Tukey test on inter-population diversity mean differences between treatment categories (n = 3).

| | cycle 4 | | cycle 10 | |
|---|---|---|---|---|
| **contrast** | **t** | **adj. p-val.** | **t** | **adj. p-val.** |
| Anc. ~ ANC* | -2.25 | 0.2390 | -3.57 | 0.0323 * |
| Antag. ~ ANC* | -2.42 | 0.1857 | 3.15 | 0.0635 ‡ |
| Cheat. ~ ANC* | 1.55 | 0.5541 | -2.73 | 0.1186 |
| no-DP ~ ANC* | -0.95 | 0.8717 | 0.28 | 0.9985 |
| Antag. ~ Anc. | -0.18 | 0.9997 | 6.72 | <0.0001 * |
| Cheat. ~ Anc. | 3.80 | 0.0277 * | 0.84 | 0.9108 |
| no-DP ~ Anc. | 1.30 | 0.6987 | 3.85 | 0.0212 * |
| Cheat. ~ Antag. | 3.98 | 0.0173 * | -5.88 | 0.0011 * |
| no-DP ~ Antag. | 1.48 | 0.5980 | -2.87 | 0.0954 ‡ |
| no-DP ~ Cheat. | -2.50 | 0.1658 | 3.00 | 0.0779 ‡ |

Significant ($p < 0.05$) and nearly significant ($0.1 > p > 0.05$) values are highlighted with * and ‡ respectively.
